# 5xFAD-NSG mice: a preclinical Alzheimer model for human cell therapy studies

**DOI:** 10.64898/2026.03.03.709048

**Authors:** David J Graber, Marie-Louise Sentman, W. James Cook, Charles L Sentman

**Affiliations:** Center for Synthetic Immunity, Department of Microbiology and Immunology, the Geisel School of Medicine at Dartmouth, Lebanon, NH 03756 USA

**Author notes:** Corresponding author: Professor Charles Sentman, Department of Microbiology & Immunology, 6W Borwell Building, One Medical Center Drive, Lebanon, NH 03756 USA. **Competing Interest Statement:** The authors (DJG, WJC, and CLS) have patents and pending patent applications for chimeric antigen receptors (CARs) and the use of CAR T-regulatory cells as therapy for neurodegenerative diseases. DJG and CLS are founders of Black Bear Bio, LLC. These interests are managed through the policies of Dartmouth College. **Funding:** This work was supported in part by grants from the National Institutes of Health AG067971, AG081632, AG090275 and funds from the Center for Synthetic Immunity, the Geisel School of Medicine at Dartmouth.

**Keywords:** Alzheimer’s disease, human T regulatory cell, mouse model, neuroinflammation, neurodegeneration, cell therapy, amyloid-beta

## Abstract

Cellular therapies are one class of medicine being developed to treat Alzheimer’s disease (AD), a neurodegenerative disease with memory loss, aberrant protein accumulation in the brain, and neuroinflammation. One challenge for development of human cell therapeutics is to have preclinical animal models that allow the use of human cells without immune-mediated rejection of those cells. The 5xFAD transgenic mouse model is a robust disease model for AD that expresses human amyloid precursor protein and presenilin-1 with multiple familial AD mutations, develops microglia-mediated neuroinflammation, accumulates amyloid-beta (Aβ) in the brain, and shows behavior changes with age. To create a mouse model for AD that permits the transplantation of human cells without immune-mediated rejection, we bred the 5xFAD transgenes onto the NOD-SCID-IL-2Rγ-deficient (NSG) mouse model to create 5xFAD-NSG mice. We report that 5xFAD-NSG mice develop Aβ plaques in the brain, microgliosis, neuroinflammation, and behavior changes that increase with age. We demonstrate that injection of CAR engineered human T regulatory cells are detected in the cerebral cortex and spleen after eight days. This 5xFAD-NSG mouse model for AD will be helpful to develop human cell-based therapies for AD.

**Significance Statement:** 5xFAD transgenic NSG mice develop a progressive AD-like disease and are an *in vivo* model for testing human cell-based therapies for AD.

## Introduction

Cell therapies are the transfer of autologous or allogeneic stem cell- and non–stem cell-living drugs to treat patients for a variety of diseases (Long & Holtzman, 2019). One recent FDA-approved example is YESCARTA®, autologous engineered T cells that target tumor cells for the treatment of relapsed or refractory large B-cell lymphoma. Stem cell- and T cell-based therapies are now being developed for the treatment of neurodegenerative diseases, which have few treatments and with limited efficacy.

Alzheimer’s disease (AD) is a progressive neurodegenerative disease that is one of the primary reasons for memory dysfunction and dementia after 60 years of age. Neuronal dysfunction and death in the frontal cortex and hippocampus, along with microglia-mediated neuroinflammation and formation of aberrant amyloid-beta (Aβ) peptide aggregates and intraneuronal tangles of twisted tau protein fibrils are hallmarks of AD (Long & Holtzman, 2019). Advancing age increases AD prevalence, with an estimated 14 million individuals in 2060 expected to be living with clinical AD in the United States (Rajan et al., 2021). Genes such as ε4 allele of the APOE gene have been identified to increase the risk of developing AD and account for 70% of the cases, whereas inherited dominant mutations in three genes have been identified that cause AD and account for an estimated range of 1.7% to 11.6% of cases (Nicolas et al., 2024),

Developing therapeutics to stop or even slow AD disease progression have proven difficult. To gain further insights into disease pathology and therapeutic interventions, several transgenic (Tg) murine models of AD have been developed by expressing one or more of the three human genes that carry a known autosomal dominant AD mutation (Zhong et al., 2024). The three genes are amyloid precursor protein (APP), presenilin-1 (PSEN1), and presenilin-2 (PSEN2). Aβ peptide, which forms aggregates and plaques in AD, is cleaved from the extracellular part of APP by PSEN1 and PSEN2 enzymes. The Tg2576 Tg mouse model overexpresses human APP with the Swedish double mutation (K670N/M671L). The APP/PS1 Tg model combines the Swedish mutant human APP gene with a PSEN1 delEx9 mutation. The PS2APP Tg model expresses the Swedish mutant human APP gene and the human PSEN2 gene containing the N141I mutation. 5xFAD Tg mice express the human APP gene with Swedish, Florida (I716V), and London (V717I) mutations, and human PSEN1 with both the M146L and L286V mutations. These different Tg models vary on the ages of onset and severity for the development of human Aβ plaque formation, neuroinflammation, and memory impairments (Yokoyama et al., 2022). AD Tg mouse models have been useful in preclinical studies for studying small molecule drugs, antibody-based therapies, and antisense oligomers.

Cell-based therapies such as with regulatory T cells (Tregs) or stem cells have been garnering interest for AD and other neurodegenerative diseases. The ability to directly study human cell therapies, as opposed to using corresponding mouse cells, in conventional AD murine models is a challenge due to the host-versus-graft human cell rejection by the murine immune system. NOD-SCID-IL-2Rγ-deficient (NSG) mice have been developed to allow the transplantation and engraftment of human cells due to their lack of T, B, and NK cells. NOG mice are similar to NSG, but rather than having a deletion of IL-2Rγ it produces a non-signaling IL-2 receptor. Furthermore, the SIRPα expressed in NSG and NOG mice naturally recognizes the human CD47 receptor as ‘self’, thus murine cells receive a ‘do not eat me’ signal that prevents mouse macrophages from attacking human cells (Ito et al., 2002; Yamauchi et al., 2013), which makes it advantageous over other immunodeficient mouse models. NSG and NOG mice permit the transplantation of human cells, tissues, and tumors and have been widely used for the development of human CAR T cell therapies for cancer (Mhaidly & Verhoeyen, 2020).

AD models have been developed on immunodeficient C57BL/6 mice that still have strong microglia-mediated neuroinflammation (Mancuso et al., 2024; Marsh et al., 2016) which is one of several attractive therapeutic targets. Recently, single and double knock-in mice were developed carrying the human APP Swedish double mutation (K670N/M671L) and human PSEN1^M146V^ in NOG mice (Yeapuri et al., 2025). In the current study, we have bred the 5xFAD AD genotype (human APP gene with Swedish, Florida (I716V), and London (V717I) mutations, and human PSEN1 with both the M146L and L286V mutations) onto the NSG background as an additional AD mouse model that permits testing of human cell therapies *in vivo*. We describe the disease development and phenotype of these 5xFAD-NSG mice, and demonstrate that systemically injected human Tregs are detected in the spleen and cerebral cortex.

## Results and Discussion

### Generation of 5xFAD-NSG mice

To create an *in vivo* model for AD that would allow the testing of human therapeutic cells, we bred 5xFAD mice to NSG mice to create a 5xFAD-NSG mouse strain. In brief, male 5xFAD mice were bred to female NSG mice, and the resulting male F1 progeny were backcrossed to NSG mice. Male offspring were used to breed with female NSG mice because the IL-2Rγ chain is found on the X chromosome. The BC1 male mice were typed for the human 5xFAD transgenes by PCR, then blood was taken to type for the absence of lymphocytes by flow cytometry. About 25% of the BC1 mice were both 5xFAD and had no T cells in the blood. We chose these male mice to bred to NSG mice to create the BC2 progeny. The male offspring were typed for the 5xFAD transgene and for a lack of blood CD3+CD4+ T cells by flow cytometry to confirm their phenotype. Male mice from BC2 were selected and bred to NSG female mice to create the BC3 progeny. All 5xFAD transgenic BC3 male mice had a white coat color and lacked blood T cells. We took BC3 male mice and bred them to NSG females to create an expanded cohort of BC4 mice for experimental use and analysis for amyloid-beta plaques and evidence for neuroinflammation. We have continued to take male offspring typed for the 5xFAD transgene for breeding of more than 14 generations. We refer to this mouse strain as 5xFAD-NSG mice. The survival of 5xFAD-NSG mice was similar to age-matched non-transgenic littermates (NTLs) through 12 months.

### Progressive Amyloid-beta plaque development in the brain of 5xFAD-NSG mice

The five mutations in the human APP and PS1 transgenes used to create 5xFAD mice results in the development of AD-like Aβ plaque formation in the brain of immune intact mice (Forner et al., 2021; Oakley et al., 2006). To assess the development of Aβ plaques in 5xFAD-NSG mice, we collected brain samples from mice at various ages from 13 to 26 weeks. As shown in Figure 1A, immunohistochemistry for Aβ was performed using Aβ recombinant rabbit monoclonal antibody (H31L21) that recognizes both human and mouse Aβ. In non-Tg littermates (NTL), diffuse background staining and darker nuclei staining were observed. Prominent Aβ plaques were only found in 5xFAD-NSG brains, and they were detected by 13 weeks of age. Amyloid plaques were observed in both male and female mice. The plaques were readily identifiable in the isocortex, subiculum and thalamus regions, and less so in the striatum. The number of amyloid plaques and the percent of plaque area in the isocortex and hippocampus increased further from 13 to 26 weeks of age, as shown in Figure 1B & 1C.

**Figure 1.**
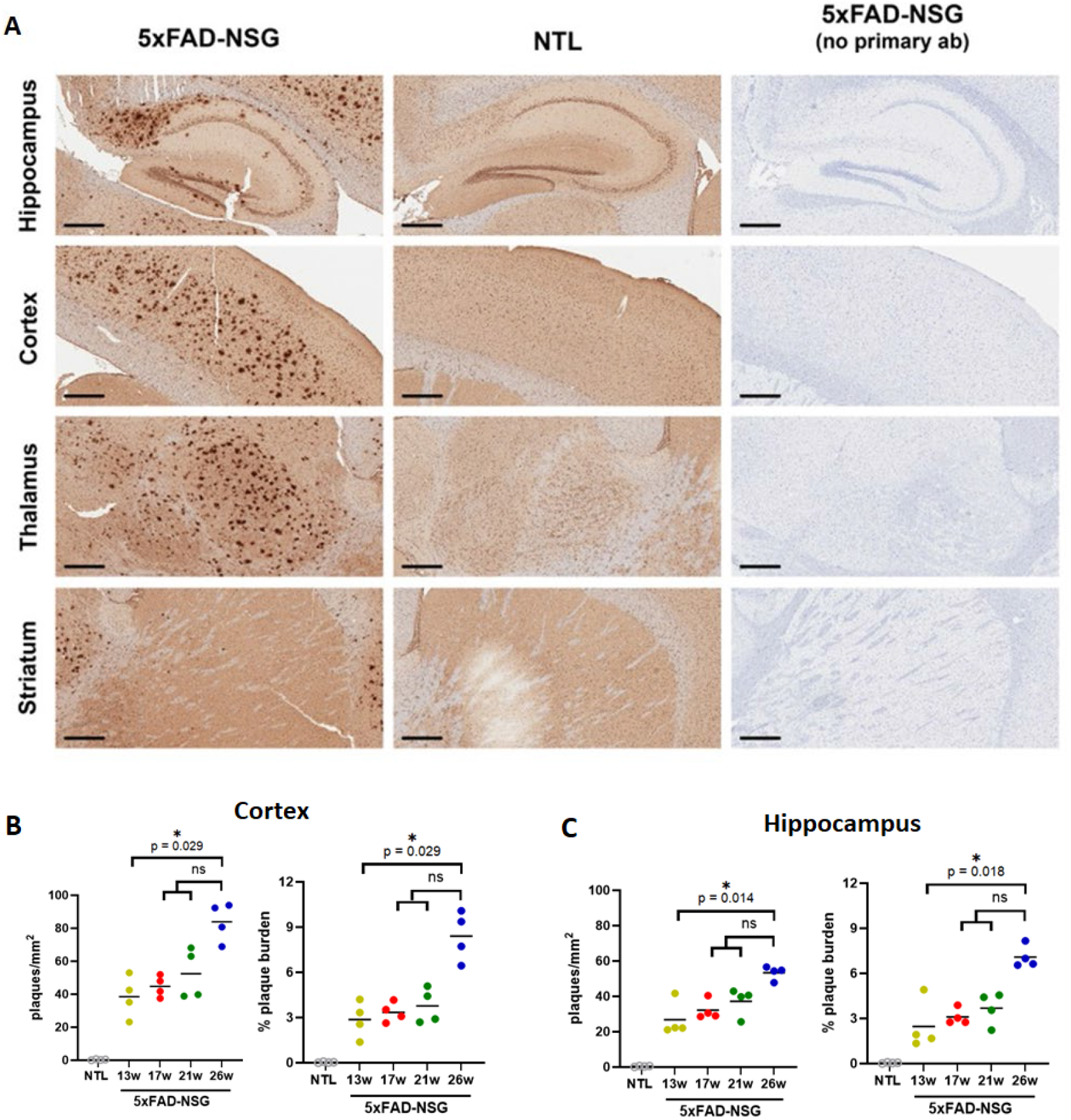
Human amyloid plaques develop in 5xFAD-NSG mouse brains. A. Representative immunohistochemical staining of anti-beta-Amyloid antibody showing plaque formation in different brain regions from 26-week aged 5xFAD-NSG mouse and no plaques observed in non-Tg littermate (NTL). Panels on the right are staining controls omitting the anti-beta-Amyloid antibody. Scales each represent 400 μm. B. Quantification of plaques per area or percent plaque burden by age in the cerebral cortex shows more plaques in 5xFAD-NSG mice at 26-week relative to 13-weeks. C. Quantification of plaques per area or percent plaque burden in the hippocampus shows more plaques at 26-week relative to 13-weeks. n = 4 per 5xFAD-NSG age group, median shown by horizonal line, p value determined by Dunn’s multiple comparison test among 5xFAD-NSG ages, ns means not significant. Four NTLs were a combined with one mouse from each age group.

### Activated microglia increase in the brain of 5xFAD-NSG mice

Activation of microglia is a prominent feature of AD disease in humans (Fan et al., 2017; Malvaso et al., 2023) and the 5xFAD transgenic mouse model (Forner et al., 2021). To evaluate how microglia activation changed over time on the NSG mouse strain, immunohistochemistry for Iba1 expression was done on brain tissues from non-transgenic littermates (NTL) and 5xFAD-NSG mice sacrificed at 13, 21, or 35 weeks of age. As shown in Figure 2A,B, the Iba1 staining is enhanced relative to the NTLs. There was prominent Iba1 microglia in the isocortex, subiculum, and the thalamus region which correspond to where Aβ plaques were prominent. There was increased Iba1+ microglia in the striatum and dentate gyrus although fewer plaques found in these areas. The percent of Iba1+ cells increased from 3.5% at 13 weeks in the total forebrain tissue up to about 10% of the total tissue area by week 35 (Figure 2C). The isocortex showed greater amounts of Iba1, which also increased over time from 5% of the cortex at 13 weeks up to 14% at 35 weeks of age (Figure 2D). There was significantly more Iba1+ microglia relative to NTLs by 21 weeks of age.

**Figure 2.**
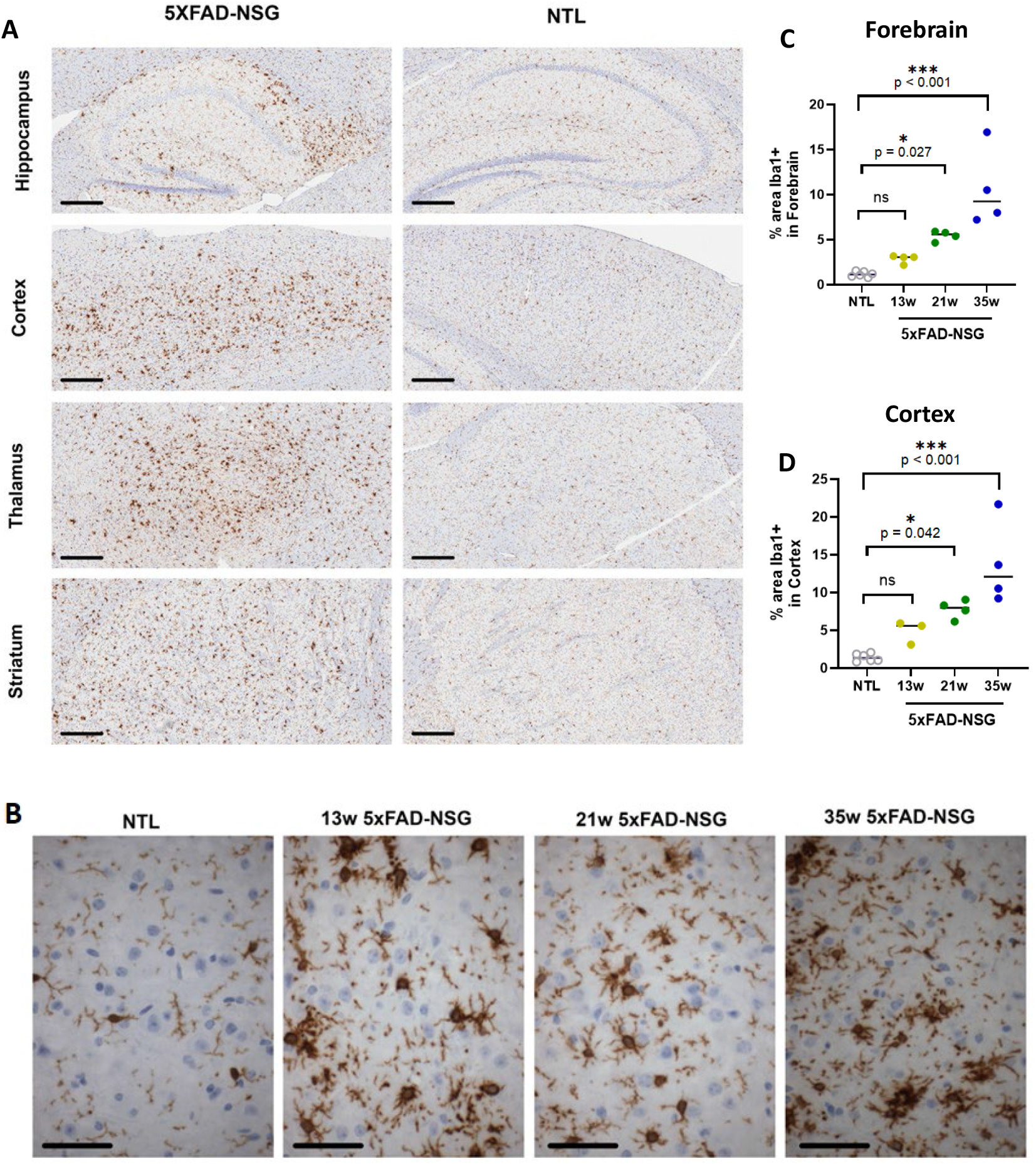
Microglia activity increases with age in 5xFAD-NSG mouse brains. A. Representative immunohistochemical (IHC) staining of Iba1 antibody showing increased microglia in different brain regions from 21-week old 5xFAD-NSG mouse relative to microglia observed in non-Tg littermate (NTL). Scales each represent 300 μm. B. Higher magnification IHC images of Iba1 antibody showing microglia morphology changes in 5xFAD-NSG mice at 13, 21, and 35 weeks of age. Scales each represent 50 μm. C. Quantification of the extent of Iba1-labeled microglia by age in the forebrain shows microglia are increased in 5xFAD-NSG mice at 13-(n = 4), 21-(4), and 26-week (4) relative to NTL (6). D. Quantification of the extent of Iba1-labeled microglia by age in the cerebral cortex shows microglia are increased in 5xFAD-NSG mice at 13-(n = 3), 21-(4), and 26-week (4) relative to NTL (6). Median shown by horizonal line, p value determined by Dunn’s multiple comparison test 5xFAD-NSG ages relative to NTL. The NTL group was combined from 2 mice from each age group. ns means not significant.

### Prominent neuroinflammation gene expression in the cerebral cortex

Neuroinflammation is a hallmark of AD and is one target for therapeutic intervention in AD. Because Aβ plaques are prominent and microglia showed activation in the 5xFAD-NSG mice, we evaluated the mRNA expression of eight inflammatory mediators known to be increased in 5xFAD B6 mice. Fresh tissue was isolated from the cerebral cortex containing both the isocortex and hippocampus regions, RNA isolated and cDNA prepared. By 13 weeks of age, there was already an increase in CCL4, Cst7, Clec7a, and Itgax expression relative to age-matched NTLs, while increase in TNF-α, IL-6, IL-1β, and NOX2 expression occurred starting around 26 weeks (Figure 3A). It is known that the disease progression differs between men and women in AD (Kolahchi et al., 2024), and female 5xFAD mice develop disease at an earlier time than male 5xFAD mice (Forner et al., 2021; Sil et al., 2022). To assess differences between male and female mice, we compared gene expression and found no difference in the time course or increased expression of TNF-α or NOX2 (Figure 3B). However, at 21-weeks of age female 5xFAD-NSG mice had higher mRNA expression of CCL4, Cst7, Clec7a, Itgax, IL-6, and IL-1β than males. This female increase was often transient with males even having higher amounts of IL1β and CCL4 at older ages. Transient sex differences in microglia activation have been reported on the B6 background (Forner et al., 2021).

**Figure 3.**
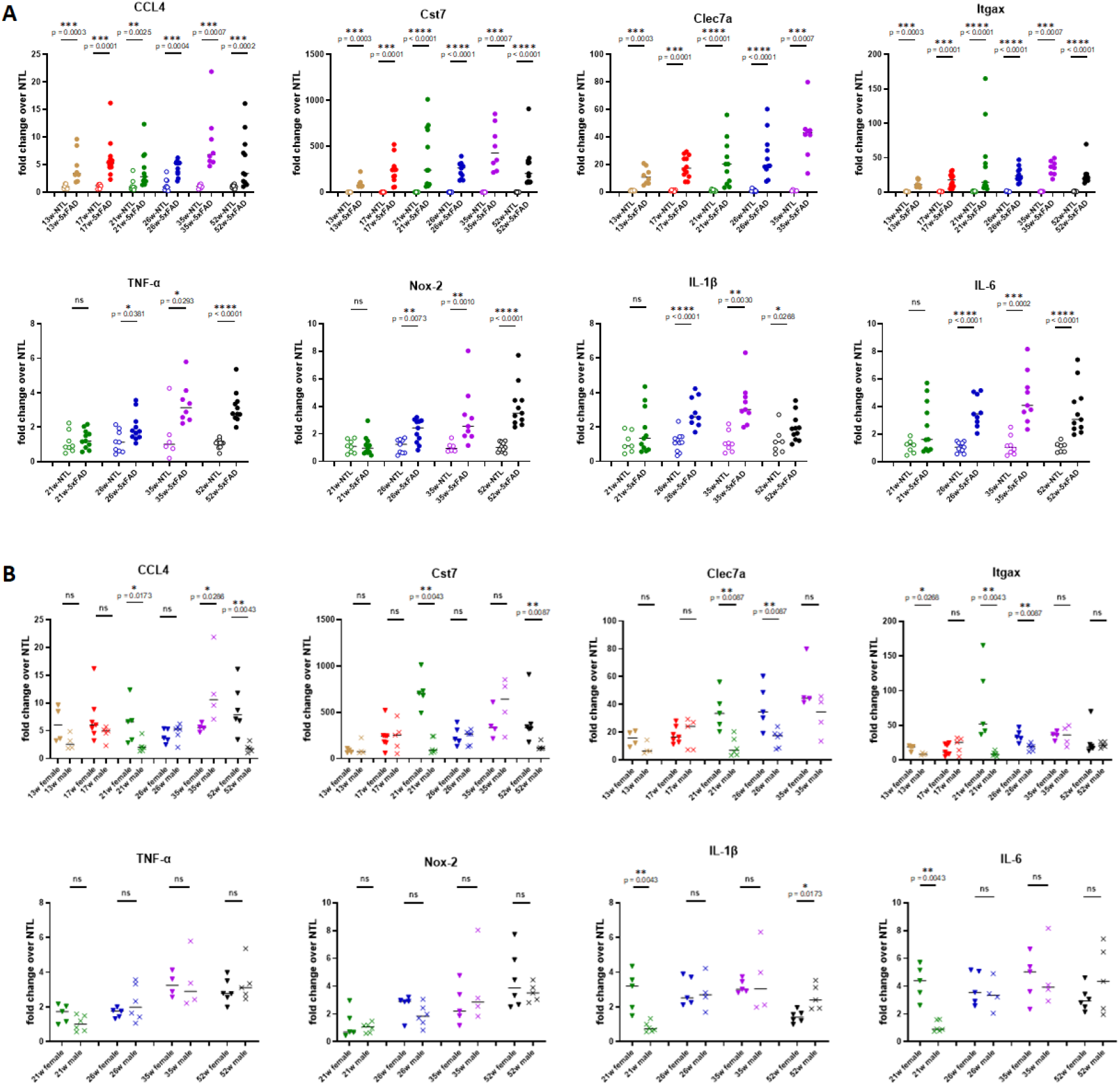
Inflammatory mediator expression changes in cerebral cortex of 5xFAD-NSG mice at different ages. A. CCL4, Cst7, Clec7a, Itgax, TNF-α, Nox-2, IL-1β, and IL-6 mRNA increased by 13 weeks or 26 weeks of age in collective male and female 5xFAD-NSG mice relative to age-matched NSG non-Tg littermate (NTL). B. Female 5xFAD-NSF mice had age-dependent higher mRNA expression than males in each inflammatory mediator except TNF-α and Nox-2, while males had higher age-dependent expression only in CCL4 and IL-1β. Values are expressed as fold-change over values from age-matched NTLs. p values determined by unpaired Mann Whitney test. * p < 0.05; ** p < 0.01; *** p < 0.001; **** p < 0.0001. ns means not significant.

### Behavior testing of 5xFAD-NSG mice using the EPM

To evaluate the behavior differences between 5xFAD-NSG and NTL mice, each mouse was tested using an elevated platform maze (EPM). The percent of time spent in the center, open arms or closed arms of the maze was evaluated for each mouse. Both male and female mice were evaluated at 23, 29 and 44 weeks of age. The 5xFAD-NSG mice spent a greater amount of time in the open arms of the maze and less in the closed arms compared to the NTL mice at 44 weeks of age (Figure 4). There were no significant differences at 23 and 29 weeks of age, although there were trends for the 5xFAD-NSG mice to spend less time in the closed arms of the maze. These data are consistent with studies using the 5xFAD-B6 strain that also show that transgenic mice spent a larger amount of time in the open arms and less in the closed arms compared to NTL littermates (Forner et al., 2021).

**Figure 4.**
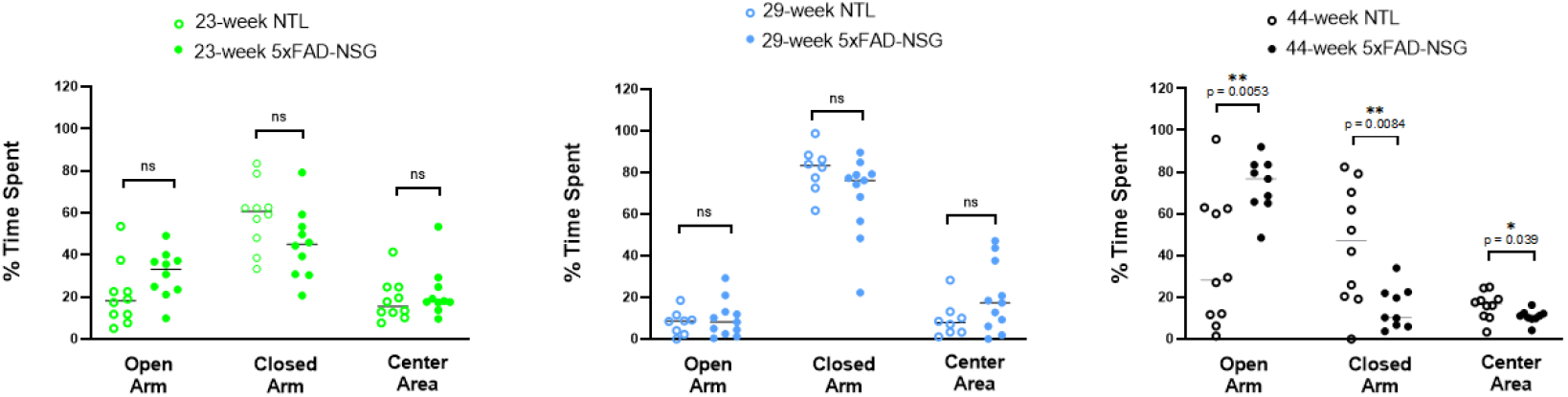
Behavior in 5xFAD-NSG mice at different ages tested in the elevated plus maze test of anxiety. Increased time spent in the open arm, an indication of reduced anxiety, was observed in 5xFAD-NSG mice at 44 weeks of age relative to age-matched non-Tg littermates (NTL). p values determined by unpaired t-test with Welch’s correction. * p < 0.05; ** p < 0.01. ns means not significant.

### Systemic injection of human Tregs expressing an Aβ-specific CAR results in human T cells in the brain and spleen

The aim of creating the 5xFAD-NSG model is to allow transplantation of human cells without rejection from the mouse immune system. In a similarly developed mSOD1-NSG model of amyotrophic lateral sclerosis, injected human Tregs persisted for many weeks whereas human Tregs were rapidly eliminated and undetectable within 2 days post injection when immunocompetent mice were used (Graber et al., 2021). To test whether human T cells can be transplanted systemically and be detected in tissues, we produced Aβ specific chimeric antigen receptor (CAR) Tregs that were expanded *in vitro* for 12 or 17 days. These CAR Tregs co-expressed a truncated mouse CD19 marker that was detected in CAR-transduced, but not mock-transduced, human Tregs (Figure 5A). The *in vitro*-expanded human Tregs maintained FoxP3 and CTLA4 expression. The Aβ-specific CARs designed with a CD44 co-stimulatory domain (BA38) or a conventional CD28 co-stimulatory domain (DG03) both secreted IL-10 *in vitro* in response to Aβ-coated wells (Figure 5B). After intravenous (IV) injection of these human CAR Tregs, the 5xFAD-NSG mice were given human IL-2 by intraperitoneal (IP) injection on days 2, 4, and 6 to support Treg survival *in vivo*. After 8 days, mice were euthanized and spleen and cortex tissues harvested. The presence of human Tregs was determined using a qPCR analysis of human CD52 mRNA, a gene that is highly expressed in human Tregs and is not expressed in mice (Graber et al., 2025). As shown in Figure 5C, human CAR Tregs were detected in the spleen (91% of all CAR Treg injected mice) and cortex (65% of all CAR Treg injected mice) of mice 8 days after injection. The relative CAR Tregs amounts in the spleen were higher than in the cortex. This 5xFAD-NSG mouse model allows the survival and tracking of human cells after IV injection, and human Aβ CAR Tregs can be detected in CNS tissue.

**Figure 5.**
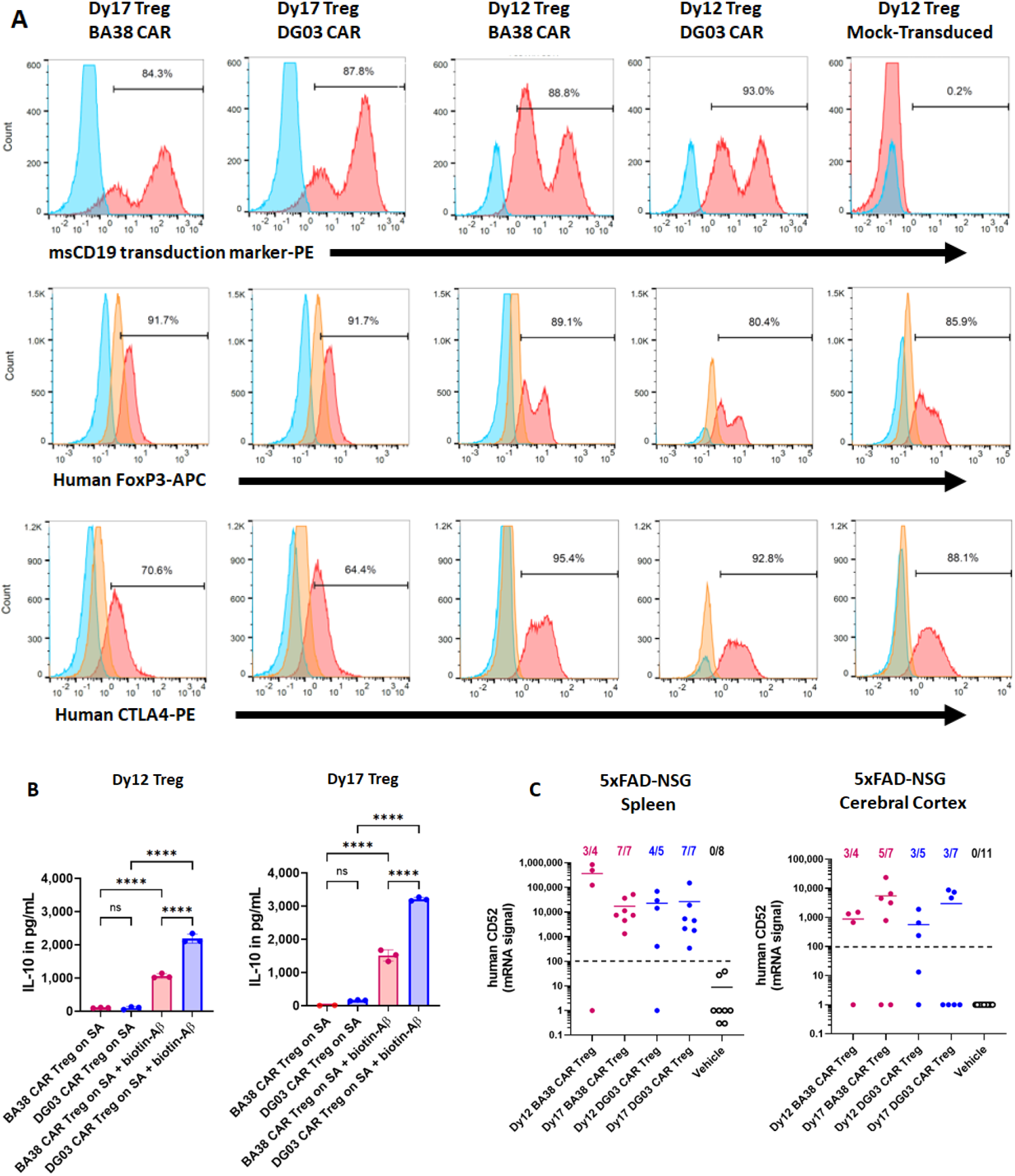
Human Tregs expressing Aβ-specific CARs are detected in 5xFAD-NSG mice after systemic injection. A. Human Tregs isolated from PBMCs, expanded *in vitro* for 12 or 17 days, and transduced to express anti-Aβ CARs with CD3ζ stimulatory domain and with either a CD44 (BA38 CAR) or a CD28 (DG03 CAR) co-stimulatory domain are shown by flow cytometry to express CAR co-transduction marker truncated mouse (ms) CD19, and Treg intracellular markers FoxP3 and CTLA4. Red is stained, blue is non-stained, and orange is isotype controls for intracellular staining. Mock-transduced Tregs were used as negative control for CAR transduction staining. B. CAR Tregs expanded for 12 or 17 days secreted IL-10 *in vitro* after plating on biotinylated Aβ bound to plate-bound streptavidin (SA), but not SA alone. p values determined by Šídák’s multiple comparisons test. N = 3 per group. *** p < 0.0001. ns means not significant. C. Eight days following intravenous injection of human CAR Tregs in 5xFAD-NSG mice, detection of human CD52 mRNA (defined as greater than 100-fold-change over the assay detection limit = 1) was observed in spleens and cerebral cortex of most mice, but not in mice injected with vehicle. Circles represent individual mice and horizontal lines represent median.

### Limitations of the 5xFAD-NSG model

While the 5XFAD-NSG mice are a valuable preclinical AD model to evaluate human cell therapies, there are several limitations to take into consideration. NSG mice the lack host T, B, and NK cells. One consequence of this lack of lymphocytes is that there is more space available for the expansion of transplanted human cells and no competition between transplanted and host lymphoid cells. There is also no source of IL-2 from host lymphocytes, which is needed to help support the survival of transplanted human Tregs *in vivo*. This is also a particular challenge in the CNS, even in immune intact animals, in which IL-2 levels are exceptionally low under normal conditions (Yshii et al., 2022). In addition, due to the lack of the IL-2Rgamma chain in NSG mice, the mice cannot respond to the cytokines that use this receptor chain, such as the type I IL-4 receptor. There has been a report of mild gliosis and vacuolation foci in the CNS of some NSG mice, and an even higher frequency and severity in NOG mice, that is predominantly found in the pons and medulla oblongada (Finesso et al., 2023). We have not observed obvious signs of neuroinflammation in the cortex, striatum, thalamus, or hippocampus in the NTLs of our 5xFAD-NSG mice or in spinal cord tissue in the NTLs of the mSOD1-NSG model (Graber et al., 2021).

Another potential limitation is that the human and murine receptor-ligand interactions may be incompatible or suboptimal. While cross-species activities are compatible for several cytokines (e.g. G-CSF, TNF-α, FLT3, TGF-β), others are known to not be compatible (e.g. IL-3, M-CSF, GM-CSF) (Hess et al., 2021). The compatibility of other factors is not fully understood (e.g. LFA-1, VLA-4, ICAM-1, VCAM-1, PD-1). This may affect the full therapeutic activity of the human cell therapy, and trafficking of the cell therapies to target tissues may be under-represented.

### Final Conclusions

The 5xFAD-NSG mouse model provides a valuable tool for developing preclinical human cell therapies for AD. These mice develop characteristic amyloid plaques and neuroinflammation in relevant brain regions with age, and develop abnormal behavior as shown with the EPM test. We demonstrate that intravenously administered human CAR Tregs avoid rejection and can be detected in both the spleen and brain tissues. Ongoing and future work in CAR Tregs in the 5xFAD-NSG mouse model will explore the enhancement of CAR Treg persistence and disease modifying activity. The 5xFAD-NSG model with its AD features and engraftment acceptance of human cells has great potential in the development of CAR Treg and other human cell therapies (e.g. human stem cells) for AD.

## Materials and Methods

### Animals

NOD.Cg-Prkdc^scid^ IL2rγ^tm1Wjl^ (NSG) mice were purchased from the Dartmouth Mouse Modeling Shared Resource (Lebanon, NH). 5xFAD-B6 mice (MMRRC Strain #034848-JAX) were purchased from the Jackson Laboratory (Bar Harbor, ME). Mouse experiments and procedures were ethically conducted under the approval of Dartmouth College’s Institution Animal Care and Use Committee. Mice were bred and colonies maintained at Geisel School of Medicine’s Center for Comparative Medicine and Research facility (Lebanon, NH) in accordance with institutional guidelines.

### Mouse genotyping

5xFAD mice were genotyped using methods described by The Jackson Laboratory with slight modifications. Briefly, small mouse tail tips were digested overnight using proteinase K (1 mg/mL, Invitrogen 25530-049), in Tris buffer (100 mM NaCl, 10 mM EDTA, and 50 mM TrisHCl), and 1 – 1.5 μL of tail genomic DNA was tested by PCR for the FAD transgenes. Primers for the 5xFAD transgene PCR genotyping assay were 27367: CGGGCCTCTTCGCTATTAC, 37598: ACCCCCATGTCAGAGTTCCT, and 37599: TATACAACCTTGGGGGATGG and were purchased from IDT. Assay conditions were 1.30 X Kapa 2G HS buffer, 2.60 mM MgCl2, 0.26 mM dNTPs, 0.03 U/μL Kapa 2G HS taq polymerase (kit KK5512), and 0.05 μM each primer. PCR cycling settings were exactly as described by The Jackson Laboratory (https://www.jax.org/Protocol?stockNumber=008730&protocolID=31769). All 25 μL of the individual PCR reactions were visualized on 2-2.5% agarose gels.

### RNA/cDNA production & qPCR analysis

Mouse tissues including cerebral cortex and spleen were isolated and frozen in RNAprotect (Qiagen 76104) as described by the manufacturer. RNA was purified using a modified procedure incorporating initial Trizol (Invitrogen 15596026) purification followed by secondary purification over RNeasy columns (Qiagen 74104) as per both manufacturers’ protocols. One or 2.5 μg of total RNA was converted to cDNA using a cDNA synthesis kit (Quanta Perfecta 84657) following the manufacturer’s protocol. One or two μL of the resulting cDNA was used in qPCR analyses using a SYBRgreen kit according to the manufacturer (Quanta Perfecta 84069) on a BioRad CTX96 instrument. Primers were designed using a web-based program (IDT PrimerQuest); sequences are listed below. Data analysis was performed using software provided by BioRad.

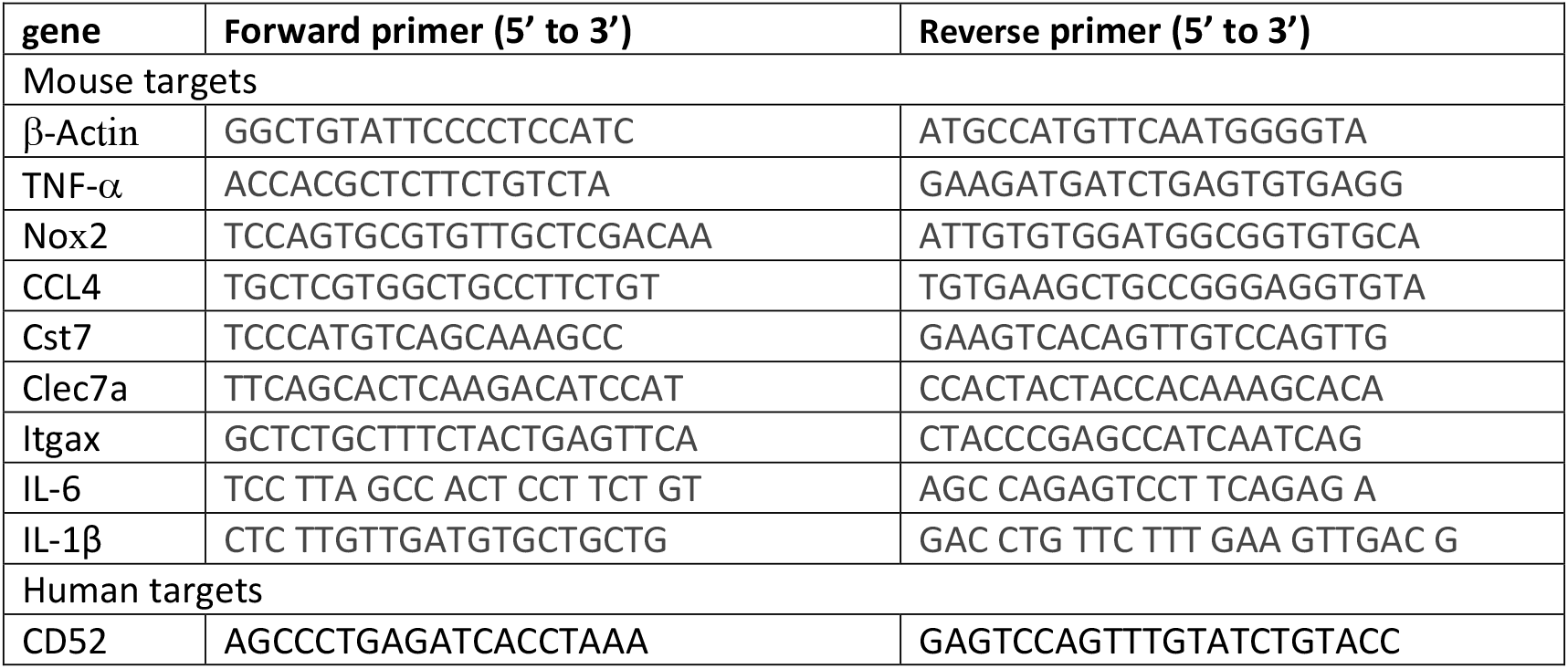

### Immunohistochemistry

Mouse tissue samples were fixed in formalin overnight, and paraffin-embedded by Pathology Shared Resource (Dartmouth-Hitchcock hospital). Tissue sections were cut at 4μm and air dried at ambient temperature before baking at 60°C for 30 min. Automated protocol performed on the Leica Bond Rx includes paraffin dewax, antigen retrieval and staining. Heat induced epitope retrieval using Bond Epitope Retrieval 2, pH9 (Leica Biosystems AR9640) was incubated at 100°C for 20-30 min. Primary antibodies were applied at room temperature (RT) with the following parameters: Beta-Amyloid antibody (Thermo 700254) incubated for 15 min at RT at a 1:100 dilution; and Iba1 antibody (Cell Signaling 17198) incubated for 15 min at a 1:500 dilution. Primary antibody binding was detected and visualized using the Leica Bond Polymer Refine Detection Kit (Leica Biosystems DS9800) with DAB chromogen and Hematoxylin counterstain. All tissue sections were digitally scanned with Leica Aperio AT2 at 20X magnification. Higher magnification (60X) images were captured using an Olympus BX51TF microscope and an Olympus Q-Fire camera. Allen Brain Atlas (mouse.brain-map.org) was used to facilitate identification of brain regions.

Iba1 expression was assessed using QuPath software (ver. 4.2). Each animal provided one or two sections per analysis and total tissue or regions of interest (ROI) were selected. Analysis settings were: resolution high (2.2 mm/px), channel DAB, prefilter Gaussian, smoothing 2.0, threshold 0.2. Each section was analyzed for expression of Iba1 on the total forebrain, isocortex, or hippocampal regions, and the average area measured per slide was 3.8 × 10^7^ mm^2^, 8.7 × 10^6^ mm^2^, and 3.4 × 10^6^ mm^2^, respectively. The total % of area that was Iba1+ was reported.

Amyloid-β plaques were assessed using ImageJ software (ver. 1.53e). Each animal provided two sections per analysis. TIFF image files were captured at 4X magnification with Aperio ImageScope software and then converted from RGB to 8-bit in ImageJ. The image threshold range was set to 0 and 64, and the particle parameters were size of 30 to infinity and circularity 0.1 to 1.0. An average of 6 mm^2^ of hippocampus and 24 mm^2^ of cortex was analyzed per mouse. Number of detected particles per mm^2^ and % of area were calculated in the isocortex or hippocampal regions.

### Elevated Platform Maze (EPM)

Mouse behavior was evaluated using an EPM for 5 min. Each mouse was gently placed in the center of the maze. The location of the mice was recorded on a video camera positioned in a holder 5 feet above the center of the maze. Data were evaluated using EthoVision software (Noldus) to determine the percent of time that each mouse spent in the center, open arms, or closed arms of the EPM. The maze was wiped down with 70% ethanol between mice and allowed to air dry for 5 min.

### Isolation of human Tregs

The protocol for enriching, sorting, expanding, and transducing human Tregs has been described in detail (Graber et al., 2025). In brief, human CD4+ cells were purified using a human CD4+ cell negative selection RosetteSep reagent as directed by the manufacturer (Stem Cell #15022, Vancouver, BC, Canada) from human blood cones obtained from plateletpheresis. CD4+ T cells were stored frozen in liquid N_2_ and then thawed in cold XVI-VO-15 media, and then rinsed in cold MojoSort Buffer (Biolegend 480017, San Diego, CA, USA). Cells were labeled with anti-CD25-PE (1/50, Biolegend 356103) and then with anti-PE magnetic beads (Miltenyi 130-105-639, Bergisch Gladbach, Germany), followed by selection over an MS column (Miltenyi 130-042-401). CD25-enriched CD4 T cells were stained with Zombie UV (Biolegend 423107) in PBS and then CD4-FITC (1/100, Biolegend 317408) and CD127-APC (1/50 Biolegend 351316). Purified human Tregs were isolated by cell sorting based on Zombie UV neg., CD4+, CD25hi, and CD127lo cells using a FACSAria-II cell sorter. After sorting, cells were counted and stimulated with CD3-CD28 tetramer antibody complexes (StemCell 10971) in XVIVO-15 media with 10% heat-inactivated human AB serum (Sigma H3667, St. Louis, MO, USA), and 500 IU/mL human IL-2. After 9 days, cells were restimulated with CD3-CD28 tetramer complexes and then spin-transduced with DG03 or BA38 CAR PG13 retrovirus on Retronectin-coated 24-well non-tissue culture plates on days 10 and 11. Cells were returned to XVIVO-15 with human sera and IL2 on day 12 and expanded until day 17. Cells on day 12 or 17 were characterized and injected into mice.

### DG03 and BA38 anti-Aβ CAR Retroviral Vector Production

Retroviruses were constructed on a pSFG backbone. Retroviral CAR vectors encoding the Aβ targeting CAR was made by fusing anti-Aβ crenezumab scFv sequences to gene fragments from CD28 hinge, transmembrane, a CD28 or CD44 costimulation domain, and the CD3ζ signaling domain. The CAR with a CD28 co-stimulatory domain is called DG03. The CAR with a CD44 co-stimulatory domain is called BA38. CARs co-expressed a non-active truncated mouse CD19 gene separated by a T2A self-cleaving peptide. Amino acid sequences were converted to DNA using the IDT’s online codon optimization software tool (www.idtdna.com/CodonOpt, Commercial Park, Coralville, Iowa, USA), which also removes unwanted restriction sites and hairpin regions in the mRNA. Restriction sites used for cloning were added to the 3′’ and 5′’ ends. Gene blocks were then supplied by IDT. Ecotropic MuLV stocks were generated by co-transfecting the retroviral vector and pPsiEco (Clontech 53460, Mountain View, CA, USA) into 293T cells. The 293T virus supernatants were harvested two days after transfection and filtered through a 0.45 -μm filter (Millex SLHVR33RS, Duluth, GA, USA) before immediate use or storing at −80°C. These ecotropic viruses were used to transduce PG13 (GALV envelope) packaging cells to generate virus to infect human cells. PG13-packaged virus supernatants were harvested from confluent cultures and filtered through a 0.45 μm filter, and storing stored at −80°C. Virus-producing cell lines 293T and PG13 were cultured in complete DMEM with a high glucose concentration (Cytiva SH30022, Marlborough, MA, USA) supplemented with 10% heat-inactivated FBS (GE HyClone SH30910, Logan, UT, USA), 100 U/mL penicillin/100 mg/mL streptomycin (Hyclone SV30010, Logan, UT, USA), 1 mmol/L sodium pyruvate (Corning Cellgro 25000-CI), 10 mmol/L HEPES (Corning Cellgro 25060-CI, Corning, NY, USA), 0.1 mmol/L MEM non-essential amino acids (Corning Cellgro 25025-CI), and 50 μM 2-mercaptoethanol (Sigma M3148).

### In vivo injection of human CAR Tregs

Female transgenic 5xFAD-NSG mice at age 25 or 32 weeks were IV injected with 8-10 × 10^6^ transduced human Tregs that were rinsed twice in PBS and resuspended in an injection solution consisting of 0.4 mL X-VIVO-15 media with IL2 (10,000 IU/mL) without serum. Mice were injected with IL-2 in 0.4 mL PBS IP at two (40,000 IU/mouse), four (10,000 IU/mouse), and six (10,000 IU/mouse) days after injection of human Tregs. Vehicle-injected mice also received IL-2 IP injections. Eight days after injection the cerebral cortex and spleen were harvested in RNA protect solution for measurement of human CD52 mRNA.

### Flow cytometry of human CAR Tregs

Human CAR Tregs on the day of injection were harvested, counted, washed twice and stained with mAbs, as indicated, in flow buffer (PBS 1% FBS). Cells are stained with mAbs at 4°C in the dark for 20min, washed, and put in flow buffer for analysis. Cells are gated on live cells using FSC and SSC. Flow cytometry antibodies (Biolegend): mouse CD19-PE, human CD4-APC, human FoxP3-APC, and FoxP3 isotype-APC, human CTLA4-PE, and CTLA4 isotype-PE. Intracellular FoxP3 staining followed membrane permeabilization using eBioscience™ Foxp3 / Transcription Factor Staining Buffer Set (Invitrogen). Cell staining was analyzed with a MACSQuant flow cytometer (Miltenyi) and FloJo v10.8.1 software.

### Human CAR Treg Cytokine Stimulation Assay

For CAR-stimulated cytokine production, cell-free cell culture media was collected after Tregs at 24h after treatment with plate-bound biotinylated Aβ_1-42_. Streptavidin (5 μg/mL, 50 μL/well, Biolegend) in carbonate buffer (Biolegend) was coated on ELISA plate (Corning) for 2 h at RT, rinsed with PBS (Corning Cellgro 21040-CV) four times and wells were then blocked with 5% FBS in PBS for 30 min. Aβ_1-42_ biotinylated at the C-terminus (200 nM, 50 μL/well AnaSpec AS-61484-01) in 5% FBS was added for 2 h at 37C, then rinsed with PBS four times. CAR Tregs were added at 50,000 cells/well in 200 μL of cell culture media without IL2. Human IL-10 was measured in the cell-free medium by ELISA using kits (BioLegend) according to the manufacturer’s directions.

### Statistics

Statistical analysis was conducted using GraphPad Prism Software (La Jolla, CA, USA). Significant differences were defined as *P < 0.05, ** P < 0.01, *** P < 0.001, and **** P < 0.0001. Data were assessed for normality using Prism software. If a Gaussian distribution was present, data were analyzed using a Student’s t-test with Welch’s correction or multiple group comparisons were done by one-way ANOVA and Šídák’s multiple comparisons test. If the data did not have a normal distributed, data were evaluated using a Mann-Whitney U test for two groups or a Kruskal-Wallis ANOVA for multiple group comparisons.

## Acknowledgements

The authors thank the staff of the Center for Comparative Medicine and Research for support with animal work, and the staff of the Department of Pathology at Dartmouth Hitchcock Medical Center for tissue preparation and IHC (RRID: SCR_023479). The authors thank Gary Ward for cell sorting, and Radu Stan for help with imaging (RRID: SCR_025077). Cell sorting was carried out in DartLab, the Immune Monitoring and Flow Cytometry Shared Resource (RRID: SCR_019165) at the Norris Cotton Cancer Center at Dartmouth, and the Genomics and Molecular Biology Core with sequencing (RRID: SCR_021293 and S10OD030242), with a National Cancer Institute Cancer Center Support Grant 5P30 CA023108. The authors also thank the National Cancer Institute Biological Resources Branch for the recombinant human IL-2. Finally, the authors thank Dr. Linda Van Eldik, University of Kentucky, for advice related to AD mouse models.

## References

Fan, Z., Brooks, D. J., Okello, A., & Edison, P. (2017). An early and late peak in microglial activation in Alzheimer’s disease trajectory. Brain, 140(3), 792–803.

Finesso, G., Willis, E., Tarrant, J. C., Lanza, M., Sprengers, J., Verrelle, J., Banerjee, E., Hermans, E., Assenmacher, C.-A., & Radaelli, E. (2023). Spontaneous early-onset neurodegeneration in the brainstem and spinal cord of NSG, NOG, and NXG mice. Veterinary Pathology, 60(3), 374–383.

Forner, S., Kawauchi, S., Balderrama-Gutierrez, G., Kramár, E. A., Matheos, D. P., Phan, J., Javonillo, D. I., Tran, K. M., Hingco, E., & Da Cunha, C. (2021). Systematic phenotyping and characterization of the 5xFAD mouse model of Alzheimer’s disease. Scientific Data, 8(1), 270.

Graber, D. J., Cook, W. J., Sentman, M.-L., Murad-Mabaera, J. M., Stommel, E. W., & Sentman, C. L. (2025). Human CAR Tregs Targeting SOD1 and Expressing BDNF Reduce Inflammation and Delay Disease in G93A hSOD1-NSG Mice. Cells, 14(17), 1318.

Graber, D. J., Sentman, M.-L., Cook, W. J., & Sentman, C. L. (2021). mSOD1-NSG mice: A new in vivo model to test human T cells in ALS. bioRxiv, 2021–09.

Hess, N. J., Brown, M. E., & Capitini, C. M. (2021). GVHD pathogenesis, prevention and treatment: Lessons from humanized mouse transplant models. Frontiers in Immunology, 12, 723544.

Ito, M., Hiramatsu, H., Kobayashi, K., Suzue, K., Kawahata, M., Hioki, K., Ueyama, Y., Koyanagi, Y., Sugamura, K., & Tsuji, K. (2002). NOD/SCID/γ c null mouse: An excellent recipient mouse model for engraftment of human cells. Blood, The Journal of the American Society of Hematology, 100(9), 3175–3182.

Kolahchi, Z., Henkel, N., Eladawi, M. A., Villarreal, E. C., Kandimalla, P., Lundh, A., McCullumsmith, R. E., & Cuevas, E. (2024). Sex and gender differences in Alzheimer’s disease: Genetic, hormonal, and inflammation impacts. International Journal of Molecular Sciences, 25(15), 8485.

Long, J. M., & Holtzman, D. M. (2019). Alzheimer disease: An update on pathobiology and treatment strategies. Cell, 179(2), 312–339.

Malvaso, A., Gatti, A., Negro, G., Calatozzolo, C., Medici, V., & Poloni, T. E. (2023). Microglial senescence and activation in healthy aging and Alzheimer’s disease: Systematic review and neuropathological scoring. Cells, 12(24), 2824.

Mancuso, R., Fattorelli, N., Martinez-Muriana, A., Davis, E., Wolfs, L., Van Den Daele, J., Geric, I., Premereur, J., Polanco, P., & Bijnens, B. (2024). Xenografted human microglia display diverse transcriptomic states in response to Alzheimer’s disease-related amyloid-β pathology. Nature Neuroscience, 27(5), 886–900.

Marsh, S. E., Abud, E. M., Lakatos, A., Karimzadeh, A., Yeung, S. T., Davtyan, H., Fote, G. M., Lau, L., Weinger, J. G., & Lane, T. E. (2016). The adaptive immune system restrains Alzheimer’s disease pathogenesis by modulating microglial function. Proceedings of the National Academy of Sciences, 113(9), E1316–E1325.

Mhaidly, R., & Verhoeyen, E. (2020). Humanized mice are precious tools for preclinical evaluation of CAR T and CAR NK cell therapies. Cancers, 12(7), 1915.

Nicolas, G., Zaréa, A., Lacour, M., Quenez, O., Rousseau, S., Richard, A.-C., Bonnevalle, A., Schramm, C., Olaso, R., & Sandron, F. (2024). Assessment of Mendelian and risk-factor genes in Alzheimer disease: A prospective nationwide clinical utility study and recommendations for genetic screening. Genetics in Medicine, 26(5), 101082.

Oakley, H., Cole, S. L., Logan, S., Maus, E., Shao, P., Craft, J., Guillozet-Bongaarts, A., Ohno, M., Disterhoft, J., & Van Eldik, L. (2006). Intraneuronal β-amyloid aggregates, neurodegeneration, and neuron loss in transgenic mice with five familial Alzheimer’s disease mutations: Potential factors in amyloid plaque formation. Journal of Neuroscience, 26(40), 10129–10140.

Rajan, K. B., Weuve, J., Barnes, L. L., McAninch, E. A., Wilson, R. S., & Evans, D. A. (2021). Population estimate of people with clinical Alzheimer’s disease and mild cognitive impairment in the United States (2020–2060). Alzheimer’s & Dementia, 17(12), 1966–1975.

Sil, A., Erfani, A., Lamb, N., Copland, R., Riedel, G., & Platt, B. (2022). Sex differences in behavior and molecular pathology in the 5XFAD model. Journal of Alzheimer’s Disease, 85(2), 755–778.

Yamauchi, T., Takenaka, K., Urata, S., Shima, T., Kikushige, Y., Tokuyama, T., Iwamoto, C., Nishihara, M., Iwasaki, H., & Miyamoto, T. (2013). Polymorphic Sirpa is the genetic determinant for NOD-based mouse lines to achieve efficient human cell engraftment. Blood, The Journal of the American Society of Hematology, 121(8), 1316–1325.

Yeapuri, P., Machhi, J., Foster, E. G., Kadry, R., Bhattarai, S., Lu, Y., Sil, S., Sapkota, R., Srivastava, S., & Kumar, M. (2025). Amyloid precursor protein and presenilin-1 knock-in immunodeficient mice exhibit intraneuronal Aβ pathology, microgliosis, and extensive neuronal loss. Alzheimer’s & Dementia, 21(4), e70084.

Yokoyama, M., Kobayashi, H., Tatsumi, L., & Tomita, T. (2022). Mouse models of Alzheimer’s disease. Frontiers in Molecular Neuroscience, 15, 912995.

Yshii, L., Pasciuto, E., Bielefeld, P., Mascali, L., Lemaitre, P., Marino, M., Dooley, J., Kouser, L., Verschoren, S., & Lagou, V. (2022). Astrocyte-targeted gene delivery of interleukin 2 specifically increases brain-resident regulatory T cell numbers and protects against pathological neuroinflammation. Nature Immunology, 23(6), 878–891.

Zhong, M. Z., Peng, T., Duarte, M. L., Wang, M., & Cai, D. (2024). Updates on mouse models of Alzheimer’s disease. Molecular Neurodegeneration, 19(1), 23.

